# Few-Shot Classification of Cryo-EM Micrographs Using Triplet Loss Embeddings

**DOI:** 10.1101/2025.10.01.679860

**Authors:** Alexander Ho, Brandon Sung, Sukyeong Lee, Francis T.F. Tsai

## Abstract

Micrograph quality assessment in cryo-electron microscopy (cryo-EM) presents a significant challenge: users must either manually screen thousands of micrographs or expend substantial computational resources processing potentially low-quality data. While few-shot learning has been applied to particle picking and subtomogram classification in cryo-EM, its application to micrograph-level quality assessment remains largely unexplored. We present a framework combining few-shot learning with cryo-EM micrograph classification using triplet loss embeddings. By combining real-space and Fourier-space information in an embedding network trained with triplet loss, we achieve competitive performance across multiple EMPIAR datasets using as few as 1-5 labeled examples per class. Our approach demonstrates improvements over traditional cross-entropy training, which often collapses to predicting only the majority class in the few-shot regime. These results suggest a practical framework for rapid adaptation of automated micrograph screening to new experimental conditions with minimal manual labeling.

## 1 Introduction

Cryo-electron microscopy (cryo-EM) has revolutionized structural biology by enabling near-atomic-resolution visualization of macromolecular complexes in their native states [1]. However, the technique’s success heavily depends on the quality of micrographs collected during data acquisition [2]. Poor-quality micrographs must be identified and filtered out early in the processing pipeline [3].

While recent deep learning advances have enabled automated micrograph quality assessment [4, 5], these approaches typically require substantial labeled training data and are tailored to specific quality classes, which may limit their flexibility when encountering new experimental conditions such as different detectors, grid types, or sample preparation methods. The predefined classification schema inherent in traditional supervised approaches may not easily accommodate the evolving landscape of cryo-EM instrumentation and experimental protocols.

Few-shot learning offers a promising solution by enabling models to learn from very limited labeled examples. However, applying few-shot techniques to micrograph classification presents unique bchallenges due to the high dimensionality and subtle quality indicators in cryo-EM data. While few-shot learning has been successfully applied to cryo-EM particle picking and 3D subtomogram classification, its application to 2D micrograph quality assessment has not been explored. To address these challenges, we leverage triplet loss [6], a metric learning objective that learns discriminative embeddings by optimizing relative similarity comparisons, potentially enabling models to avoid collapse to the majority class that can affect standard cross-entropy approaches in few-shot regimes.

**Figure 1.**
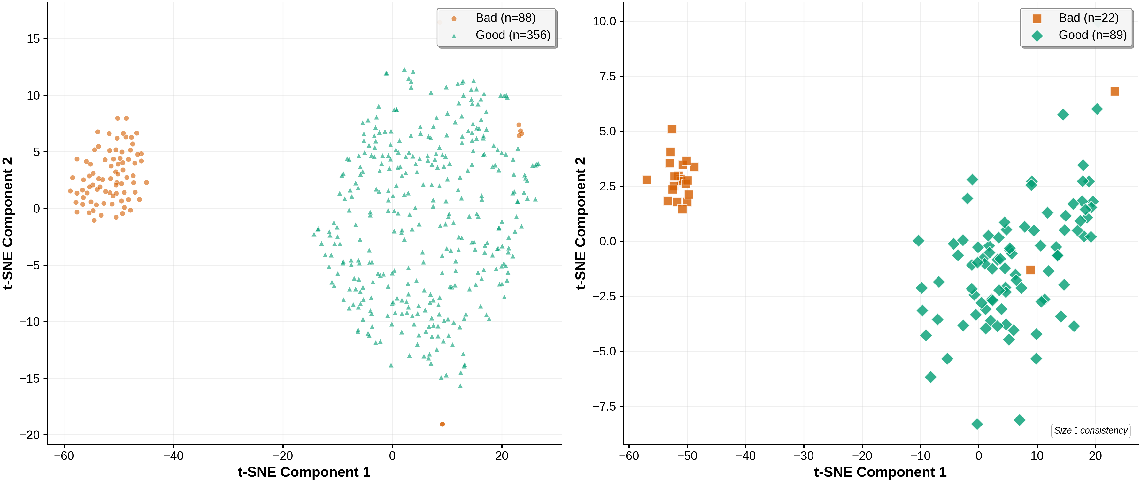
t-SNE [7] visualization of learned embeddings for hold-out test set of EMPIAR-10379 in the one-shot regime. Left: crop-level embeddings (n=88 bad, n=356 good); Right: micrograph-level embeddings obtained by averaging crop embeddings (n=22 bad, n=89 good). Clear class separation is maintained even with minimal training data.

In this work, we present a few-shot learning framework for cryo-EM micrograph classification through: (1) a dual-input architecture leveraging both real-space and Fourier-space information, (2) triplet loss training that learns discriminative embeddings to address majority class collapse, and (3) hierarchical classification aggregating crop-level predictions for robust micrograph-level decisions. We demonstrate promising results across three diverse EMPIAR datasets, achieving competitive performance with as few as 1-5 labeled examples per class.

## 2 Related Work

Traditional approaches to micrograph quality assessment have evolved from manual screening to automated deep learning methods. Recent developments include Miffi [4], which pioneered the integration of Fourier space information with CNN-based filtering, and MicAssess [5], which enables user-free preprocessing pipelines. Automated workflows like TranSPHIRE [8] and Smart Leginon [9] have incorporated deep learning for real-time quality control, while specialized tools like MicrographCleaner [10] provide targeted cleaning capabilities. However, these systems typically require substantial labeled training data, which may limit their adaptability to new experimental conditions.

The broader cryo-EM field has witnessed rapid adoption of deep learning across the processing pipeline. In particle picking, methods have evolved from early approaches like DeepPicker [11] to sophisticated tools including SPHIRE-crYOLO [12] (YOLO-based detection), Topaz [13] (positive-unlabeled learning), and more recent advances like DeepCryoPicker [14] and PIXER [15]. For image processing and reconstruction, specialized tools include Warp [16] and Topaz-Denoise [17] for denoising, CryoDRGN [18] for modeling structural heterogeneity, and DeepEMhancer [19] for map enhancement.

While few-shot learning has emerged as a promising solution for data-limited scenarios, its application in cryo-EM has been primarily focused on 3D analysis. In cryo-ET, Yu et al. [20] developed a deep Brownian distance covariance method for subtomogram classification, while Li et al. [21] demonstrated the effectiveness of Prototypical Networks. For particle picking, recent work includes CryoMAE [22], which employs masked autoencoders with self-cross similarity loss, and SaSi [23], which utilizes self-augmentation strategies. However, few-shot approaches for 2D micrograph quality assessment have not been explored.

**Figure 2.**
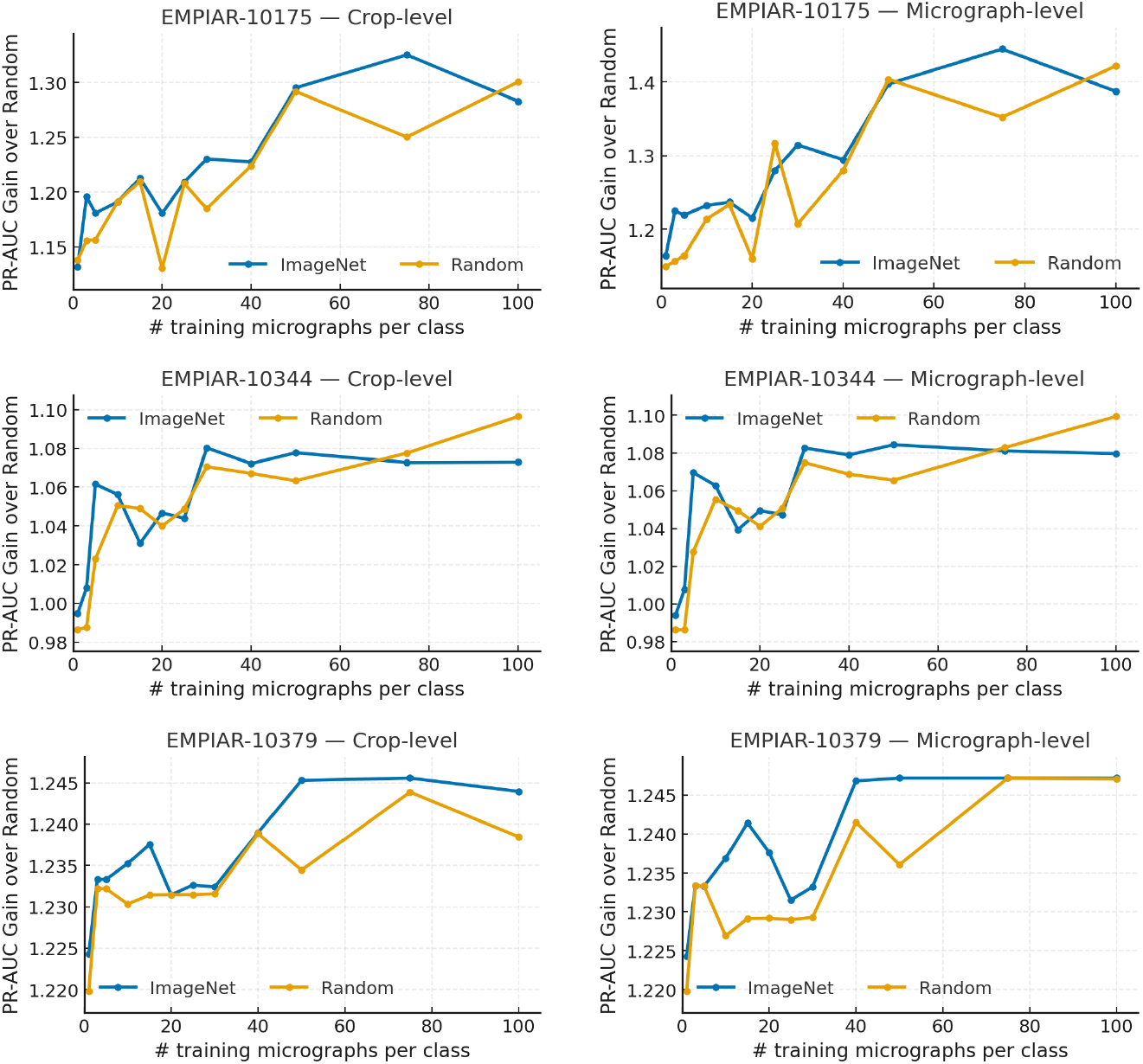
AUPRC Gain over majority class selection versus number of training micrographs per class for EMPIAR datasets 10175 (top), 10344 (middle), and 10379 (bottom). Left: crop-level results; Right: micrograph-level results after majority voting.

Our work addresses this gap by developing a few-shot framework for micrograph classification. We build upon the demonstrated importance of Fourier-space information [4] while introducing modern metric learning approaches to enable rapid adaptation to new experimental conditions with minimal labeling requirements.

## 3 Methods

### 3.1 Datasets and Preprocessing

We evaluated our approach using three EMPIAR datasets (IDs 10175, 10344, and 10379), all acquired with K2 Summit detectors [24, 25, 26]. Following [4], micrographs were labeled as “bad” if they exhibited support film issues, sample displacement, ice crystallinity, or contamination, and “good” otherwise (see Appendix Fig. S1 for representative examples). For each micrograph, we extracted both real-space crops and their corresponding centered log-scaled power spectra, incorporating both spatial and frequency-domain information. Note that while micrographs in figures are shown with 4x pixel binning for improved visual contrast (see Appendix Fig. S2), our model processes the original unbinned data.

### 3.2 Few-Shot Learning Framework

Our framework employs a dual-input ResNet18 architecture [27] that processes both real-space crops and their corresponding centered log-scaled power spectra, following the dual-input strategy insights from [4]. The backbone features optional ImageNet pretraining and is connected to an embedding network consisting of a 256-dimensional hidden layer with ReLU activation, dropout (0.5), and a 128-dimensional final embedding layer with batch normalization.

We initially explored standard cross-entropy training but found that it consistently collapsed to predicting only the majority class in the few-shot regime. Self-supervised pretraining approaches such as multi-task regression also did not help with the classification collapse. To address this limitation, we implemented a triplet loss approach [6] with online hard triplet mining, training with the Adam optimizer at a learning rate of 0.0001. For inference, we perform k-nearest neighbors classification on the learned embeddings, applying majority voting across crops to make robust micrograph-level decisions. The k parameter for k-NN classification was selected based on validation set performance.

### 3.3 Experimental Design and Evaluation

We implemented rigorous micrograph-level data splits to prevent leakage, with each crop inheriting its parent micrograph’s label. Training sets varied from 1 to 100 example micrographs per class, while validation and test sets remained fixed. We employed three random seeds for all experiments and maintained consistent data splits across model configurations. For ties in majority voting, we defaulted to “bad” classification to maintain conservative filtering.

We evaluate our few-shot classification approach using AUPRC normalized by the proportion of the majority class. Our binary classification task labels micrographs as “good” (positive class) or “bad” (negative class), where the class distribution is highly imbalanced. In our evaluation datasets, “good” micrographs constitute the majority class, though we note this could vary across datasets depending on the data source and collection process.

Critically, traditional classifiers on these imbalanced datasets exhibit pathological collapse behavior, predicting only the majority class for all examples. Standard accuracy metrics are unsuitable for this problem setting because a trivial classifier that always predicts the majority class achieves high accuracy (proportional to the class imbalance) while providing no discriminative value, obscuring the collapse behavior we seek to avoid.

The normalized AUPRC metric directly addresses this failure mode: the denominator (proportion of majority class examples) represents the baseline performance of a collapsed classifier, while values exceeding 1.0 indicate genuine discriminative ability. This normalization thus measures the practical utility we aim to demonstrate, specifically whether few-shot learning can maintain classifier discriminability on highly imbalanced data where standard approaches degenerate to trivial constant predictors.

## 4 Results

### 4.1 Performance Across Datasets

We evaluated our triplet-loss framework across three EMPIAR datasets, focusing on precision and AUPRC for the “good” class. Test-set class proportions were 0.520 (10175), 0.898 (10344), and 0.802 (10379), establishing different baselines for performance evaluation.

EMPIAR-10379 demonstrated exceptional separability, achieving perfect micrograph-level AUPRC (1.000) with just 50 micrographs per class using ImageNet initialization. Even in the extreme few-shot regime (1-5 micrographs per class), performance remained remarkably high with minimal difference between pretrained and randomly initialized models.

For the imbalanced EMPIAR-10344 dataset (89.8% “good” micrographs), AUPRC Gains approached the theoretical maximum. The best result achieved AUPRC *≈* 0.988 (Gain *≈* 1.099) at 100 micrographs per class, with precision(good) *≈* 0.984 and recall(good) *≈* 0.800. ImageNet initialization provided notable benefits in the few-shot regime, improving AUPRC Gain by 0.042 at 5 micrographs per class.

EMPIAR-10175, with the most balanced class distribution, showed consistent improvement with increased training data. Using ImageNet initialization, AUPRC Gain increased from 1.164 (1-shot) to 1.445 (75 micrographs per class), corresponding to precision(good) *≈* 0.646 and recall(good) *≈* 0.780.

Complete numerical results for all datasets, training regimes, and initialization strategies are provided in Tables S1–S6 in the Appendix.

### 4.2 Embedding Space Analysis

The t-SNE visualizations reveal clear class separation in the learned embedding space, even in the challenging one-shot regime. Crop-level embeddings show well-defined clusters despite inherent local variability, while micrograph-level embeddings obtained through averaging demonstrate enhanced separation with reduced intra-class scatter. This hierarchical organization validates our majority voting approach and confirms the effectiveness of our triplet loss training strategy.

## 5 Discussion

Our results suggest that triplet-loss embedding learning can help address the majority class collapse that may occur in few-shot micrograph classification. The method’s success varies across datasets, reflecting different levels of intrinsic separability: EMPIAR-10379’s strong performance with minimal training suggests that local features and Fourier transforms may provide discriminative signals for this dataset, while EMPIAR-10175’s more modest gains indicate underlying complexity that may require additional examples.

ImageNet pretraining appears to provide the most benefit in ultra-low-shot regimes (≤ 10 micrographs per class), potentially stabilizing embedding geometry when supervision is scarce. The consistent advantage of micrograph-level aggregation supports our hierarchical approach, with majority voting appearing to effectively integrate local evidence while reducing noise.

Our few-shot approach may help address adaptability challenges in automated micrograph screening. In contrast to traditional supervised models that typically require retraining for different experimental conditions, our framework appears to adapt to novel setups with reduced labeling effort. This flexibility could be valuable in the evolving cryo-EM landscape, where new detector technologies and imaging protocols continue to emerge. The ability to achieve competitive performance with just 1-5 labeled examples per class may make our approach practical for laboratories with diverse experimental conditions and limited annotation time.

## 6 Conclusion

We have presented a few-shot learning framework for cryo-EM micrograph classification that achieves competitive performance with minimal labeled data. Our triplet loss approach, combined with dual-input architecture and hierarchical prediction aggregation, appears to learn discriminative features while potentially avoiding the majority class collapse that can affect standard cross-entropy methods in few-shot regimes.

The framework’s potential practical impact lies in its adaptability: enabling effective classification with as few as 1-5 labeled examples per class may reduce manual labeling burden when encountering new experimental conditions. This could provide a pathway for laboratories to implement automated quality assessment across diverse experimental setups, potentially broadening access to advanced screening capabilities.

Future work could explore using pretrained weights from existing micrograph quality assessment models (e.g., Miffi, MicAssess) as initialization for triplet loss training, which might improve few-shot performance by leveraging domain-specific representations learned from larger datasets. Additional directions include extending the framework to multi-class quality assessment and investigating other metric learning objectives for cryo-EM applications.

## Supporting information

Supplemental Results

## 7 Acknowledgements

This work was supported in part by the National Institutes of Health grant R01-GM142143 and by the Welch Foundation grant Q-1530-20250403.

## A Supplementary Figures and Tables

**Figure S1:**
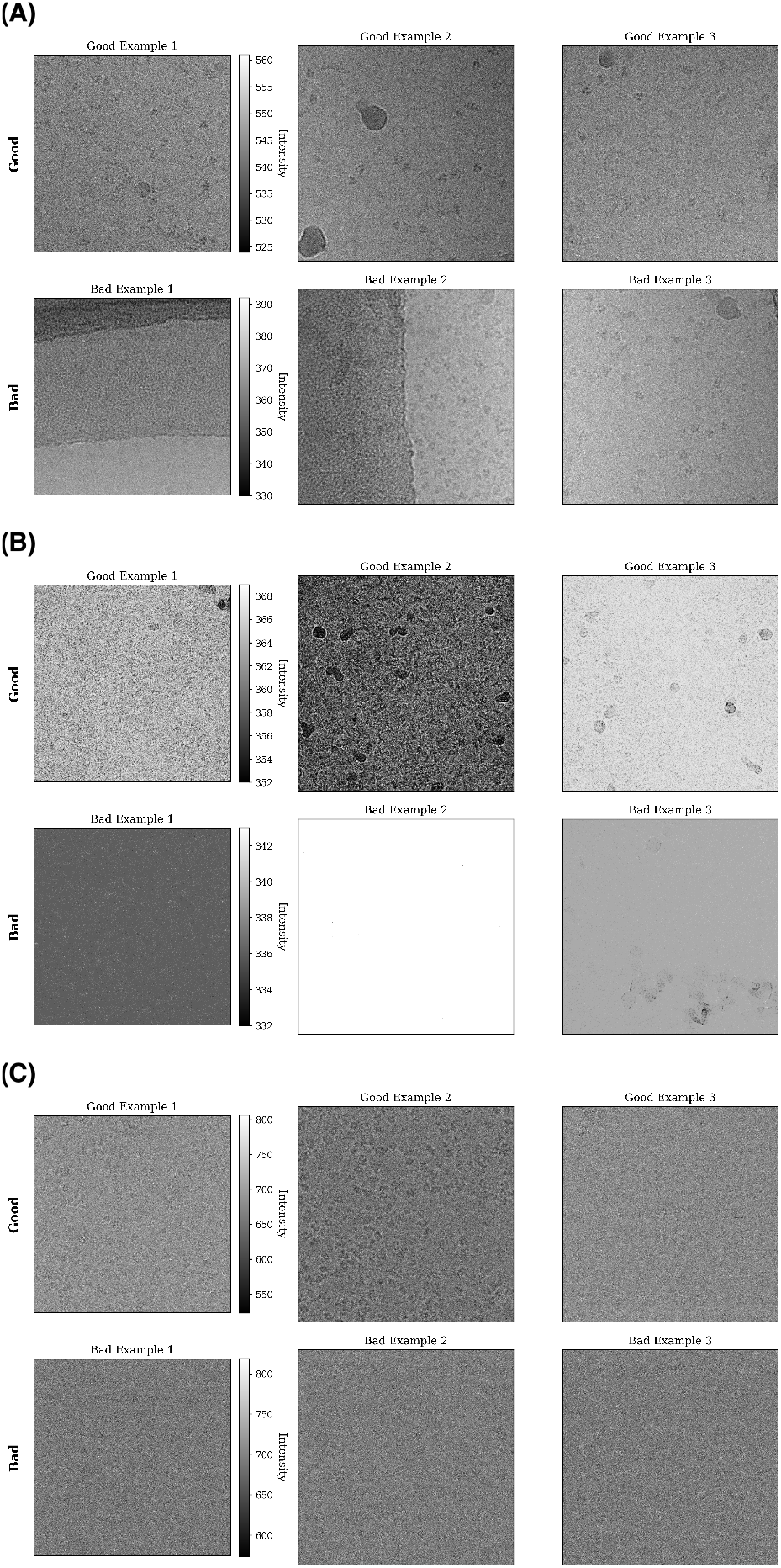
Representative examples of good and bad micrographs from EMPIAR datasets used in this study, showing the variety of defects that must be identified across different samples. For each dataset, top panels show high-quality micrographs suitable for further processing, while bottom panels show micrographs that should be rejected. Bad examples for (A) EMPIAR-10175, exhibiting support film; (B) EMPIAR-10344, showing empty and overexposed images; (C) EMPIAR-10379, showing empty images. All micrographs shown are 4x pixel-binned for improved visual contrast only.

**Figure S2:**
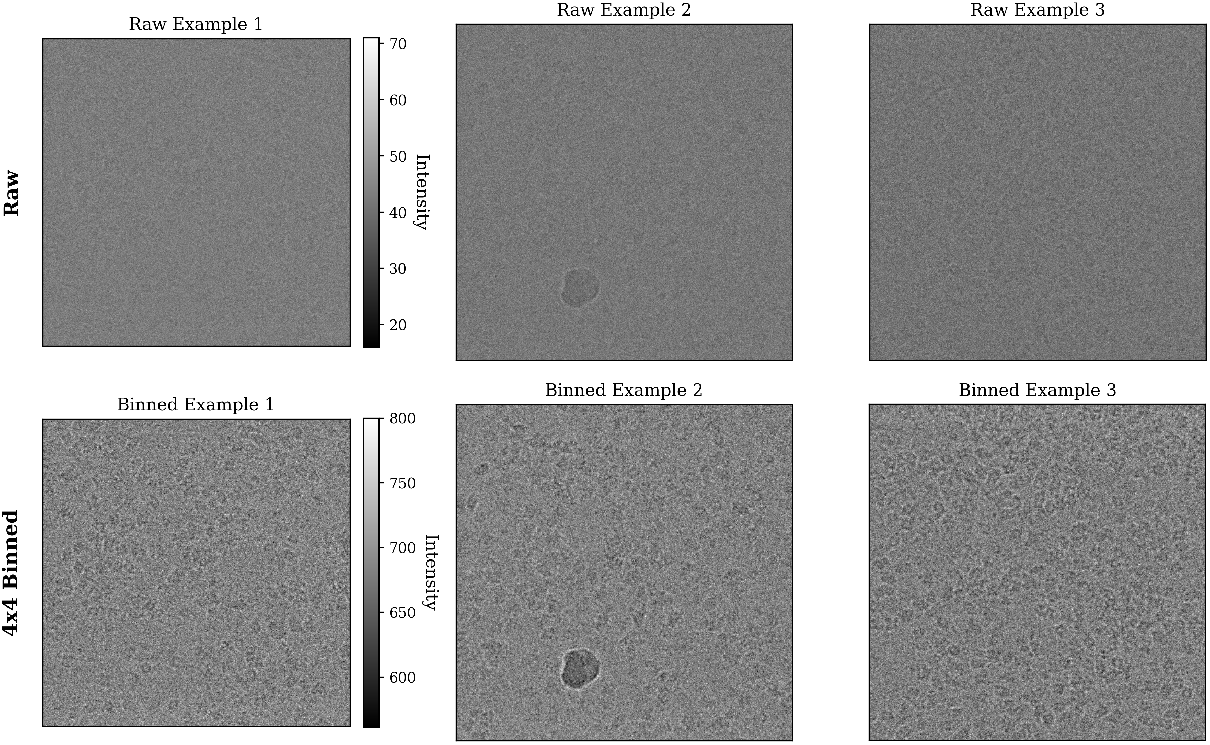
Comparison of raw and binned micrographs from EMPIAR-10379, illustrating the characteristically low SNR of cryo-EM data. Top: Original micrograph at full resolution. Bottom: The same micrograph after 4x pixel binning, showing improved contrast while preserving essential quality indicators. All micrographs shown in Fig. S1 underwent similar binning preprocessing.

**Table S1:** EMPIAR-10175 Crop-Level Results: AUPRC gain over random (AUPRC*/p*_pos_) versus number of training micrographs per class. Means and standard errors (SE) are aggregated across seeds (*N* = 3).

| Init | Shots | Mean gain | SE |
| --- | --- | --- | --- |
| ImageNet | 1 | 1.132 | 0.023 |
| ImageNet | 3 | 1.196 | 0.012 |
| ImageNet | 5 | 1.181 | 0.022 |
| ImageNet | 10 | 1.191 | 0.026 |
| ImageNet | 15 | 1.213 | 0.029 |
| ImageNet | 20 | 1.181 | 0.021 |
| ImageNet | 25 | 1.209 | 0.050 |
| ImageNet | 30 | 1.230 | 0.020 |
| ImageNet | 40 | 1.228 | 0.028 |
| ImageNet | 50 | 1.295 | 0.060 |
| ImageNet | 75 | 1.325 | 0.048 |
| ImageNet | 100 | 1.283 | 0.008 |
| Random | 1 | 1.138 | 0.001 |
| Random | 3 | 1.156 | 0.004 |
| Random | 5 | 1.156 | 0.002 |
| Random | 10 | 1.191 | 0.003 |
| Random | 15 | 1.210 | 0.012 |
| Random | 20 | 1.131 | 0.030 |
| Random | 25 | 1.208 | 0.020 |
| Random | 30 | 1.185 | 0.015 |
| Random | 40 | 1.224 | 0.041 |
| Random | 50 | 1.292 | 0.027 |
| Random | 75 | 1.250 | 0.020 |
| Random | 100 | 1.301 | 0.034 |

**Table S2:** EMPIAR-10175 Micrograph-Level Results: AUPRC gain over random (AUPRC*/p*_pos_) versus number of training micrographs per class.

| Init | Shots | Mean gain | SE |
| --- | --- | --- | --- |
| ImageNet | 1 | 1.164 | 0.028 |
| ImageNet | 3 | 1.225 | 0.011 |
| ImageNet | 5 | 1.219 | 0.026 |
| ImageNet | 10 | 1.232 | 0.047 |
| ImageNet | 15 | 1.237 | 0.035 |
| ImageNet | 20 | 1.215 | 0.051 |
| ImageNet | 25 | 1.280 | 0.070 |
| ImageNet | 30 | 1.314 | 0.031 |
| ImageNet | 40 | 1.294 | 0.052 |
| ImageNet | 50 | 1.397 | 0.094 |
| ImageNet | 75 | 1.445 | 0.066 |
| ImageNet | 100 | 1.387 | 0.032 |
| Random | 1 | 1.149 | 0.000 |
| Random | 3 | 1.157 | 0.003 |
| Random | 5 | 1.164 | 0.007 |
| Random | 10 | 1.214 | 0.003 |
| Random | 15 | 1.234 | 0.045 |
| Random | 20 | 1.160 | 0.051 |
| Random | 25 | 1.317 | 0.028 |
| Random | 30 | 1.207 | 0.019 |
| Random | 40 | 1.280 | 0.057 |
| Random | 50 | 1.404 | 0.030 |
| Random | 75 | 1.352 | 0.053 |
| Random | 100 | 1.421 | 0.046 |

**Table S3:** EMPIAR-10344 Crop-Level Results: AUPRC gain over random (AUPRC*/p*_pos_) versus number of training micrographs per class.

| Init | Shots | Mean gain | SE |
| --- | --- | --- | --- |
| ImageNet | 1 | 0.995 | 0.001 |
| ImageNet | 3 | 1.008 | 0.010 |
| ImageNet | 5 | 1.062 | 0.005 |
| ImageNet | 10 | 1.056 | 0.007 |
| ImageNet | 15 | 1.031 | 0.009 |
| ImageNet | 20 | 1.047 | 0.023 |
| ImageNet | 25 | 1.044 | 0.018 |
| ImageNet | 30 | 1.080 | 0.003 |
| ImageNet | 40 | 1.072 | 0.018 |
| ImageNet | 50 | 1.078 | 0.005 |
| ImageNet | 75 | 1.073 | 0.006 |
| ImageNet | 100 | 1.073 | 0.010 |
| Random | 1 | 0.986 | 0.002 |
| Random | 3 | 0.988 | 0.002 |
| Random | 5 | 1.023 | 0.004 |
| Random | 10 | 1.050 | 0.011 |
| Random | 15 | 1.049 | 0.004 |
| Random | 20 | 1.040 | 0.003 |
| Random | 25 | 1.049 | 0.006 |
| Random | 30 | 1.071 | 0.010 |
| Random | 40 | 1.067 | 0.003 |
| Random | 50 | 1.063 | 0.002 |
| Random | 75 | 1.078 | 0.006 |
| Random | 100 | 1.097 | 0.004 |

**Table S4:** EMPIAR-10344 Micrograph-Level Results: AUPRC gain over random (AUPRC*/p*_pos_) versus number of training micrographs per class.

| Init | Shots | Mean gain | SE |
| --- | --- | --- | --- |
| ImageNet | 1 | 0.994 | 0.000 |
| ImageNet | 3 | 1.008 | 0.012 |
| ImageNet | 5 | 1.070 | 0.006 |
| ImageNet | 10 | 1.063 | 0.007 |
| ImageNet | 15 | 1.039 | 0.011 |
| ImageNet | 20 | 1.049 | 0.024 |
| ImageNet | 25 | 1.047 | 0.020 |
| ImageNet | 30 | 1.083 | 0.003 |
| ImageNet | 40 | 1.079 | 0.019 |
| ImageNet | 50 | 1.084 | 0.006 |
| ImageNet | 75 | 1.081 | 0.005 |
| ImageNet | 100 | 1.080 | 0.012 |
| Random | 1 | 0.986 | 0.003 |
| Random | 3 | 0.986 | 0.002 |
| Random | 5 | 1.028 | 0.006 |
| Random | 10 | 1.055 | 0.011 |
| Random | 15 | 1.049 | 0.006 |
| Random | 20 | 1.041 | 0.004 |
| Random | 25 | 1.051 | 0.007 |
| Random | 30 | 1.075 | 0.009 |
| Random | 40 | 1.069 | 0.004 |
| Random | 50 | 1.066 | 0.004 |
| Random | 75 | 1.083 | 0.005 |
| Random | 100 | 1.099 | 0.003 |

**Table S5:** EMPIAR-10379 Crop-Level Results: AUPRC gain over random (AUPRC*/p*_pos_) versus number of training micrographs per class.

| Init | Shots | Mean gain | SE |
| --- | --- | --- | --- |
| ImageNet | 1 | 1.224 | 0.005 |
| ImageNet | 3 | 1.233 | 0.000 |
| ImageNet | 5 | 1.233 | 0.000 |
| ImageNet | 10 | 1.235 | 0.003 |
| ImageNet | 15 | 1.238 | 0.004 |
| ImageNet | 20 | 1.231 | 0.001 |
| ImageNet | 25 | 1.233 | 0.000 |
| ImageNet | 30 | 1.232 | 0.001 |
| ImageNet | 40 | 1.239 | 0.003 |
| ImageNet | 50 | 1.245 | 0.001 |
| ImageNet | 75 | 1.246 | 0.001 |
| ImageNet | 100 | 1.244 | 0.001 |
| Random | 1 | 1.220 | 0.000 |
| Random | 3 | 1.232 | 0.001 |
| Random | 5 | 1.232 | 0.001 |
| Random | 10 | 1.230 | 0.000 |
| Random | 15 | 1.231 | 0.001 |
| Random | 20 | 1.231 | 0.000 |
| Random | 25 | 1.231 | 0.000 |
| Random | 30 | 1.232 | 0.000 |
| Random | 40 | 1.239 | 0.003 |
| Random | 50 | 1.234 | 0.002 |
| Random | 75 | 1.244 | 0.001 |
| Random | 100 | 1.238 | 0.002 |

**Table S6:** EMPIAR-10379 Micrograph-Level Results: AUPRC gain over random (AUPRC*/p*_pos_) versus number of training micrographs per class.

| Init | Shots | Mean gain | SE |
| --- | --- | --- | --- |
| ImageNet | 1 | 1.224 | 0.005 |
| ImageNet | 3 | 1.233 | 0.000 |
| ImageNet | 5 | 1.233 | 0.000 |
| ImageNet | 10 | 1.237 | 0.005 |
| ImageNet | 15 | 1.241 | 0.005 |
| ImageNet | 20 | 1.238 | 0.004 |
| ImageNet | 25 | 1.231 | 0.000 |
| ImageNet | 30 | 1.233 | 0.000 |
| ImageNet | 40 | 1.247 | 0.000 |
| ImageNet | 50 | 1.247 | 0.000 |
| ImageNet | 75 | 1.247 | 0.000 |
| ImageNet | 100 | 1.247 | 0.000 |
| Random | 1 | 1.220 | 0.000 |
| Random | 3 | 1.233 | 0.000 |
| Random | 5 | 1.233 | 0.000 |
| Random | 10 | 1.227 | 0.000 |
| Random | 15 | 1.229 | 0.001 |
| Random | 20 | 1.229 | 0.000 |
| Random | 25 | 1.229 | 0.000 |
| Random | 30 | 1.229 | 0.001 |
| Random | 40 | 1.242 | 0.006 |
| Random | 50 | 1.236 | 0.005 |
| Random | 75 | 1.247 | 0.000 |
| Random | 100 | 1.247 | 0.000 |

Notes: Random baseline for AUPRC equals the positive-class prevalence $p_{\text{pos}}$ . Gains were computed per run as $\text{AUPRC}/p_{\text{pos}}$ using the confusion matrix to estimate $p_{\text{pos}}$ at each level, then averaged across seeds.

